# Acute stress affects peripersonal space representation

**DOI:** 10.1101/2021.04.25.441303

**Authors:** Giulia Ellena, Tommaso Bertoni, Manon Durand-Ruel, John Thoresen, Carmen Sandi, Andrea Serino

**Affiliations:** MySpace Lab, Department of Clinical Neuroscience, Centre Hospitalier Universitaire Vaudois (CHUV), University of Lausanne, Lausanne, Switzerland; Department of Psychology, University of Bologna, Bologna, Italy; CsrNC, Centre for Studies and Research in Cognitive Neuroscience, University of Bologna, Cesena, Italy; Laboratory of Behavioral Genetics, Brain Mind Institute, School of Life Sciences, Swiss Federal Institute of Technology Lausanne (EPFL), Lausanne, Switzerland; Laboratory of Cognitive Neuroscience, Brain Mind Institute, Ecole Polytechnique Federale de Lausanne, Lausanne, Switzerland

## Abstract

Peripersonal space (PPS) is the region of space surrounding the body. It has a dedicated multisensory-motor representation, whose purpose is to predict and plan interactions with the environment, and which can vary depending on environmental circumstances. Here, we investigated the effect on the PPS representation of an experimentally induced stress response. We assessed PPS representation in healthy humans, before and after a stressful manipulation, by quantifying visuotactile interactions as a function of the distance from the body, while monitoring salivary cortisol concentration. Participants, who showed a cortisol stress response, presented enhanced visuotactile integration for stimuli close to the body and reduced for far stimuli. Conversely, individuals, with a less pronounced cortisol response, showed a reduced difference in visuotactile integration between the near and the far space. In our interpretation, physiological stress resulted in a freezing-like response, where multisensory-motor resources are allocated only to the area immediately surrounding the body.

## Introduction

Peripersonal Space (PPS) is a key cortical system involved in implementing behavioural responses to environmental changes. The term PPS defines the space immediately surrounding the body, where the individual can physically interact with external stimuli in the environment. The primate brain hosts a dedicated network to represent this sector of space to detect harms to avoid, or interesting stimuli to approach when those are close to the body. PPS representation is based on a multisensory mechanism which integrates actual or even potential stimulation on the body with visual or auditory stimuli specifically presented close to the body, in a body-centred reference frame (Rizzolatti et al, 1997; Cléry et al, 2015; Graziano and Cooke, 2006; Làdavas and Serino, 2008; Serino, 2019). Original knowledge about PPS representation came from single-cell recordings in monkeys (see for a review Graziano and Cooke, 2006); neuroimaging studies in humans further revealed enhanced responses for near-body stimulation localized in posterior parietal and premotor areas of the human brain, largely corresponding to the regions where PPS neurons have been described in the monkey brain (see Grivaz, Blanke and Serino, 2017, for a review).

Those areas are directly connected, or are even part of the motor system, and, indeed direct electrical stimulation of these premotor and parietal regions hosting PPS neurons in monkeys, results in body parts movements. In humans, auditory (Serino, Annella and Avenanti, 2009; Finisguerra et al, 2015) or visual (Makin et al, 2009; Cardellicchio, Sinigaglia and Costantini, 2011) stimuli close to the body have been shown to affect the excitability of the corticospinal tract, by either inhibiting or enhancing its responsiveness, as to prepare freezing-like or active responses to stimuli within the subject’s action space. Inhibition of premotor areas (Avenanti, Annela and Serino, 2012) or parietal (Serino, Canzoneri and Avenanti, 2011) PPS regions via non-invasive brain stimulation techniques abolish these effects. Also, the hand-blink reflex, i.e., a subcortical defensive response elicited by the stimulation of the median nerve (HBR; Bisio et al, 2017; Sambo et al, 2012; Wallwork et al, 2016), is modulated as a function of the distance between the stimulated arm and the face. Thus, the multisensory representation of PPS is immediately transformed into automatic overt or potential reactions.

A wide body of evidence also shows that the extent of PPS representation is modulated as a function of the potential interactions with stimuli in the environment. PPS representation expands after tool-use (see Maravita et al, 2002, for a review), or conversely shrinks after a period of immobilization (Bassolino et al, 2010). More novel data suggest that not only sensory-motor, but also higher-level factors affect PPS representation: i.e., social interactions. PPS contracts when facing an estranger (Teneggi et al, 2013) or another person who is perceived as immoral (Iachini et al, 2015; Pellencin et al, 2018), and it extends towards other people the participants are willing to interact with (Teneggi et al, 2013; Pellencin et al, 2018). Personality traits, such as anxiety (Sambo and Iannetti, 2013), phobias (Taffou and Viaud-Delmon, 2014; Cartaud et al, 2018; Lourenco, Longo and Pathman, 2011) also contribute to defining the extent of PPS. For all these properties, PPS has been conceived as a dynamic and plastic margin around the body defining where and how the individuals potentially interact with the external environment, including the others (Cléry et al, 2015; Serino, 2019).

Stress defines a cluster of physiological and behavioural responses aimed at activating resources to face challenging situations, as well as restoring and maintaining the organism’s homeostasis (Karatsoreos and McEwen, 2011; McEwen, 2013). Depending on the species, stress-related responses are typically described by the “fly or fight” mode, which generally implies an enhanced ability to use the diverse metabolic substrates, with an increase of the blood pressure, heart rate, glycolysis, and enhanced blood supply to the central nervous and to the locomotor system to mediate a withdrawal or aggressive behaviour. A threat-related defensive strategy associated to stress is the “freezing response” (Roelofs, 2017), which is characterized by a tense body posture (reduced postural sway) with an increased muscle tone, and a reduced heart rate (bradycardia). Freezing can be defined as a state of “attentive immobility”, in which the physiological features of both sympathetic and parasympathetic are present to react to a challenging situation.

Considering the role of PPS representation in mediating environmental interactions and the impact of stress on such interactions, in the present study we asked whether and how the induction of a stress response would affect PPS representation in humans.

To this aim, we adapted a well-known experimental manipulation used to induce stress in human subjects, i.e., the Fear Factor stress test (Du Plooy et al, 2014; see Methods for a full description) and we studied its effect on PPS representation. Before and after the stress manipulation, PPS representation was measured through a multisensory task, extensively used in the literature to measure changes in PPS representation. In this task, subjects receive a tactile stimulus on their body, to which they are instructed to reply as fast as possible, while task-irrelevant auditory or visual stimuli (that have to be ignored) approach (or recede from) the body. In different conditions, tactile stimuli are delivered when the external stimuli are perceived at a different distance from the body. In line with the above-described enhancement of multisensory interaction within the PPS, several studies show that participants respond progressively more rapidly as the external stimuli approach (Canzoneri, Magosso and Serino, 2012). The relation between tactile reaction times and the position of the external stimulus in space (from here, the PPS function) is used to measure the features of PPS representation at the individual level (Ferri et al, 2015; Serino et al, 2018). To mathematically synthesize the properties of PPS representation, the relationship between tactile reaction times and distance was fitted with a linear or sigmoidal function. The central point of the fitted sigmoidal function provides a measure of the spatial position at which a looming stimulus starts to be integrated with tactile processing and affects motor reactions, thus providing an indication of the spatial boundary of PPS (Canzoneri et al, 2012; Serino et al, 2015). The slope of the linear fit quantifies the difference between the effects of near and far visual stimuli on tactile processing, providing a proxy of the amount of differentiation between peri and extrapersonal space (Noel et al, 2018; Salomon et al, 2017). Steeper PPS slopes indicate a more selective processing and motor preparation for near-body stimuli, whereas flatter slopes suggest more homogeneous monitoring of near and far stimuli with respect to potential interactions with the body.

Here, we tested how stress manipulation affects PPS representation, both in terms of its extent and its amount of differentiation. In particular, given the defensive role of PPS representation (see de Vignemont and Iannetti, 2015) and recent data about the reaction to subjectively-perceived threatening stimuli (Vagnoni, Lourenco and Longo, 2012; Taffou and Viaud-Delmon, 2014), a response to acute stress might be reflected by an extension of PPS representation, as to anticipate potential contacts with external stimuli. If this is the case, a shift towards the far space of the PPS central point should be found after stress. On the other hand, if a stress response is characterized by a “freezing” behaviour (Hagenaars, Oitzl and Roelofs, 2014; Roelofs, 2017), we would expect an enhancement of information processing in the near space, and in a reduction of resources allocated to the far space, meaning an increased steepness of the slope of the PPS function (see de Haan et al, 2016).

To verify the effectiveness of the stressor manipulation, and to measure the neuroendocrine stress response, we collected salivary samples to quantify the salivary cortisol concentration. This is normally taken as a (Hellhammer, Wüst and Kudielka, 2009) neuroendocrine measure of the ongoing stress response, operated by the hypothalamic-pituitary-adrenal axis (Hellhammer, Wüst and Kudielka, 2009). Besides, a subjective feeling of stress was also measured via a visual analog scale (VAS). Previous reports showed important individual differences in stress response, as measured by salivary cortisol changes (Kudielka, Hellhammer and Wüst, 2009). This variability is due to demographic (Kudielka and Kirschbaum, 2005), genetic (Wüst et al, 2004; Zhang et al, 2014), psychological (Iacovino, Bogdan and Oltmanns, 2016; Sandi et al, 2008; Southwick, Vythilingam and Charney, 2005) factors. Therefore, after being exposed to acute stress, some individuals may report a subjective feeling of being stressed, but no response in terms of cortisol increase, while others, may present an effective secretory episode linked to the stressing event (Campbell and Ehlert, 2012; Cohen et al, 2000; Hjortskov et al, 2004). For these reasons, here we used cortisol change to distinguish between glucocorticoid responders (*C-Responders*) and nonresponders (*Non-C-Responders*) following the exposition to the stressor (Du Plooy et al, 2014; Quaedflieg et al, 2017; Kudielka et al, 2009), as an implicit measure, and a subjective stress scale, as an explicit measure. Therefore, we compared not only PPS representation between the experimental and control group, but also within the experimental group, between responders and non-responders. Finally, both PPS representation (Sambo and Iannetti, 2013; Spaccasassi and Maravita, 2020) and stress responsiveness (Weger and Sansi, 2018; Portella et al., 2005; Everared et al, 2015; Frank et al, 2006; Roelof et al, 2010) can be influenced by individuals’ levels of anxiety. Therefore, we also investigated whether and how predisposition to anxiety, as measured by the Trait scores form of the State-Trait Anxiety Inventory (Spielberger et al, 2017) was related to both the effectiveness of the stressor manipulation and its effect on PPS representation.

## Materials and Methods

### Participants

Given the sex differences in anxiety and cortisol responsiveness (e.g., Bale and Epperson, 2015; Boettcher, Hartmann, Zimmer, and Wudy, 2017; Kudielka and Kirschbaum, 2005), only male participants were included in the present study. Thirty-eight healthy volunteers took part in the experiment (mean age 22.9±5.1years). None of the subjects reported a history of neurological or psychiatric disorders and all were naïve to the aim of the experiment. The experiment was conducted in accordance with the principles of the 1964 Declaration of Helsinki and was approved by the Ethical Committee of the Brain and Mind Institute, EPFL. Each participant gave a written informed consent prior to participating.

### Procedure

Participants were randomly assigned to one of the two groups: *Experimental* (n=19) and *Control* group (n=19). Once informed about the structure and aims of the experiment, they signed the consent form. In a first phase (the experiment timeline is illustrated in Fig. 1), that preceded the experimental manipulation (0-30 min after the start of the experiment), the stress level was measured. Two samples of saliva were collected as a physiological index of stress, and two VAS were administered, as a measure of subjective stress, in two timepoints (T1-T2). In this first phase, PPS representation was assessed via a visuo-tactile interaction task (see below for task description). The task was split into two blocks. The first was performed after the first Stress Measure (PPS-1a), and the second PPS block after the second stress measure (PPS-1b). Successively (30-80 min), the experimental group was exposed to a modified version of the Fear Factor stress test (Du Plooy et al., 2014; detailed in the next paragraph), which combines two validated tasks aimed at inducing psychophysical stress (STRESSOR1-STRESSOR2) for a sufficient amount of time. After the stress manipulation, PPS was once more assessed, again in two blocks. One block was presented after the first stressor manipulation (STRESSOR1; PPS-2a) and the other after the second (STRESSOR 2; PPS-2b). Also, cortisol and a subjective stress measure were collected in this phase (T3-T4). The Control group went through the same procedure with the exception that, instead of the experimental manipulation, two non-stressful tasks (CONTROL1-CONTROL2) were proposed (see below for control task description).

**Figure 1.**
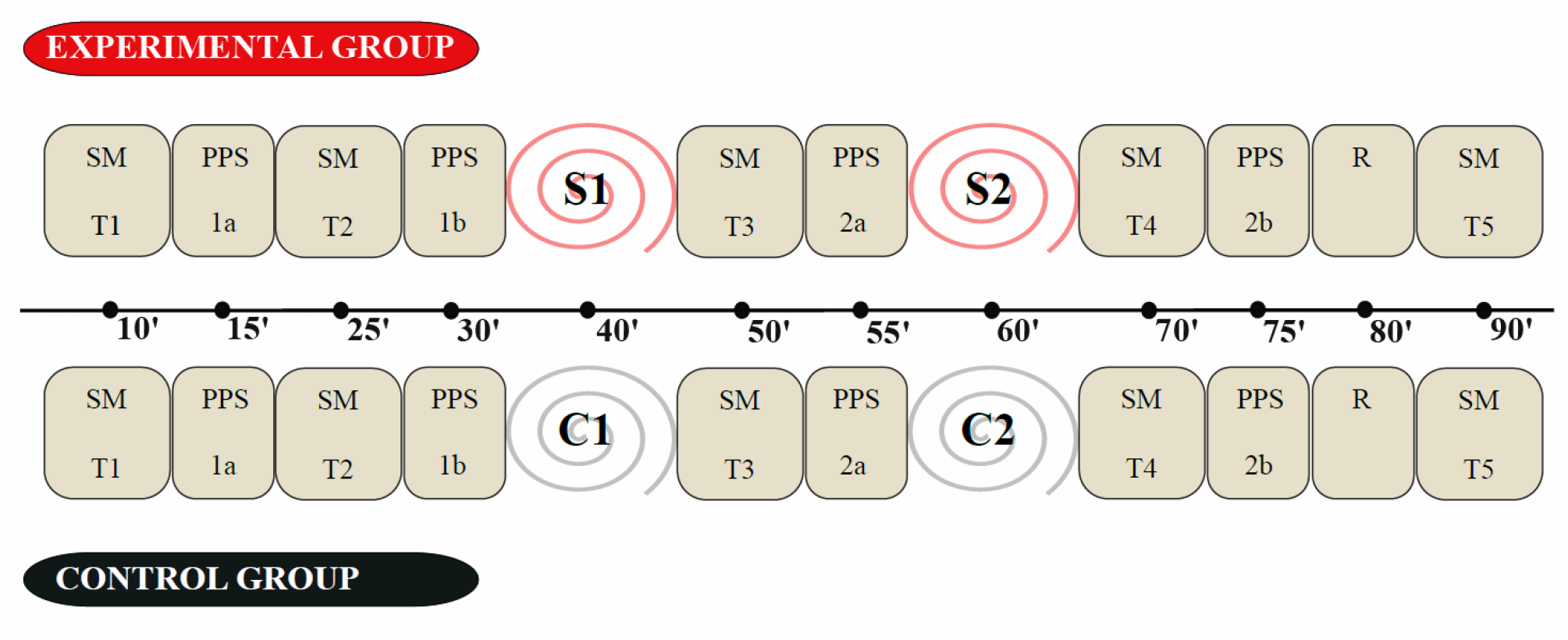
Overview of the procedure. (SM): Stress Measure. (PPS): Peripersonal Space task. (S1-S2): Stress Manipulations. (R): Release phase. (C1-C2): Control Manipulation. A between-subjects design was used: measurements form the Experimental group were compared with the Control group. The Experimental group was exposed to the stress protocol (S1+S2), while the Control group, to a non-stressing condition (C1+C2).

After the manipulation and the related stress measurements, participants underwent the final step (80-90min), the *Release* phase (R), in which they were asked to complete a self-report to assess trait anxiety and perform a distance judgment paradigm (similar to the one used for the PPS task; see below) to validate the obtained PPS measure. Finally, another cortisol sample and subjective stress measures were taken (T5). Each participant was then debriefed and paid.

### Stressor manipulation: The Fear Factor Stress Test and the control condition

The participants underwent a modified version of the Fear Factor Stress Test (Du Plooy *et al*., 2014). This stressor combines elements of the Trier Social Stress Test (Kirschbaum, Pirke and Hellhammer, 1993) and the Cold Pressor Task (CPT; Hines and Brown, 1936), the two most widely protocols used as stressors in this field of research.

Participants were asked to prepare for 2 minutes a motivational presentation to participate in a TV-reality show “Fear Factor” and then to deliver the presentation in front of a video-camera. After the motivational speech, the subjects were asked to perform a challenging arithmetic task (subtraction task) in front of the camera (STRESSOR1). The control group undertook instead a simple writing and reading task with neutral content, followed by a basic counting task instead of the subtraction. No recording occurred for the control group (CONTROL1).

The Cold Pressor task (CPT) is widely used in the psychology literature (e.g., Lovallo, 1975), and consist of the dominant arm’s immersion in cold water (~4°C) to just above the elbow, for 2 minutes. Participants from the experimental group were video-recorded during the entire water-immersion task to add a socio-evaluative component (STRESSOR2). The arm of the control participants was immersed in comfortably warm water (~35-40°C) for the same amount of time, with no video recording (CONTROL2).

### Stress measurements: Salivary cortisol levels and the Subjective Stress

To measure the level of stress, both physiological (cortisol concentration) and subjective measures were collected (STRESS MEASURE, see Fig.1).

### Cortisol Concentration Measure

To measure the cortisol level at regular time-ranges, saliva samples were taken at five time-points (Stress measure: T1-T2-T3-T4-T5; see Fig.1). A sample of approximately 0.8 to 1.4 mL of saliva was obtained at each collection in 10mL polypropylene tubes and frozen below −20 °C until processed. Samples were then centrifuged at 3000 rpm for 15 minutes at room temperature, and salivary cortisol concentrations were measured by enzyme immunoassay according to the manufacturer’s instructions (Salimetrics, Newmarket, Suffolk, United Kingdom). The samples were used to analyze cortisol baseline levels and hormonal changes taking place during the experiment. To control for the circadian rhythm of cortisol, all experimental sessions were scheduled between 1 PM and 7 PM.

#### - Assessment of Subjective Stress

To provide a measure of individual subjective evaluations in response to stress exposure, participants reported subjective ratings of stress. Subjects rated their perceived level of stress on a 1 (low) to 5 (high) visual analog scale (VAS). The measurements of the Subjective Stress were collected on five occasions (Stress measure: T1-T2-T3-T4-T5; see Fig.1).

### State-Trait Anxiety Inventory (STAI)

Participants rated their level of personal anxiety using the Trait form of the State-Trait Anxiety Inventory (STAI; Spielberger et al, 2017). Trait anxiety reflects a predisposition to anxiety as determined by the personality pattern. A French version of the STAI was used.

### Peripersonal Space Representation task

To assess PPS representation we adopted a visuo-tactile interaction task, implemented into the RealiSM software (Laboratory of Cognitive Neurosciences, EPFL) as described in Serino and colleagues (2018; see also Pellencin et al, 2018). In this task, participants are presented with task irrelevant looming visual stimuli (virtual volleyballs), while providing speeded motor responses to tactile stimuli, by pressing a button with the right hand.

The task consisted in a total of 150 trials of 5 seconds of duration, presented in different conditions, in a randomized order. In 108 trials (the 72% of the total amount) both visual and tactile stimuli were presented (visuo-tactile condition). Tactile stimuli could be delivered at a different delay from the beginning of the trial (1.82s, 2.15s, 2.475s, 2.80s, 3.12s or 3.45s), which implies that tactile information was processed when the visual stimulus was at one out of 6 distances from the participants (equally spaced from the farthest, D6 = 90 cm, to the closest D1 = 30 cm). In 24 trials (16%) only tactile stimuli were presented and no visual stimuli were shown (unimodal tactile condition). Tactile stimuli were delivered at the same six delays as for visuo-tactile stimulation. Finally, in a set of 18 trials (12%), only visual stimuli were presented (unimodal visual condition). No response was expected, and these trials were used as catch trials in order to reduce overt expectations.

The whole PPS paradigm was split into two blocks (PPS 1a/1b; PPS 2a/2b), administered as described in the procedure.

#### - Stimuli

Tactile stimuli were provided on the right jaw via a small vibrator (100 ms of duration, as in Noel et al, 2016). Visual stimuli were presented in an head-mounted display (HMD, model Oculus Rift, stereoscopic resolution 1280 x 800, diagonal field-of-view 110°), and consisted in looming volleyballs in an augmented reality scenario, where the scene background consisted of a projection of the real scene in front of the participant which was acquired by a camera (Duo3D MLX, 752 × 480 at 56 Hz) mounted on the HMD.

#### - Distance estimation task

To verify the validity of the distance manipulation, at the end of the experimental session, participants performed a distance estimation task. They were asked to estimate the distance, in meters, of the perceived looming ball position, at the different times of tactile stimulation. Distance estimation judgments provided the indication that every subject could actually discriminate six different distances (averaged values: D6 was perceived at 82 cm (SD = 14.4) far from the subject, D5 at 80 cm (SD = 6.60), D4 at 71 cm (SD = 9), D3 at 60 cm (SD = 8.5), D2 at 51 cm (SD = 9.7) and D1 at 37 cm meters (SD=9.4).

#### - PPS data analysis

In line with previous studies (Serino et al, 2015; Pellencin et al, 2018), to provide a measure of the multisensory facilitation induced by visuo-tactile stimuli on tactile processing, RT in the visuo-tactile condition were referred to the RT in unimodal tactile condition (vibration but no ball shown). For each subject, the fastest unimodal RT (after averaging per each temporal delay) was subtracted from the distance-averaged visuo-tactile RT (baseline correction RT). This correction allows estimating the multisensory gain, after controlling for a possible expectation effect due to the temporal delay of the tactile stimulation.

To obtain meaningful indices of PPS representation, the baseline-corrected RTs were fitted with a sigmoidal and a liner function to extract, respectively, the central point and the slope parameters. The central point describes the extent of PPS, whereas the slope describes the segregation of PPS from extrapersonal space (Noel et al, 2018). Units are defined so that the linear slope is expressed in the millisecond of multisensory facilitation per meter.

### Data and code availability

Behavioural data and R code for reproducing the main results are available in the following OSF repository: https://osf.io/kpdw6/?view_only=836db15416e94e51b256d524a77cde52

## Results

### Cortisol concentration

To confirm that our experimental manipulation correctly modulated stress level, we compared mean concentrations of salivary cortisol across the five measurements for the two groups (for details on saliva sampling, see Figure 1A). A mixed ANOVA with *Time* (T1/T2/T3/T4/T5) and *Group* (Experimental/Control) as factors indicated a main effect of *Time* (F(4,144)=6.35; p<0.001), of *Group* (F(1,36)=10.46; p=0.004) and a *Group* X *Time* interaction (F(4,144)=4.61; p=0.002). Newman-Keuls corrected post-hoc comparisons revealed, in the *Experimental Group*, an increase in Cortisol level from T1-T2 (which were not different from each other; p=.17) to T3 (both p-values<.001) and T4 (both p-values<.001). At T5, cortisol level then decreased to pre-manipulation levels (not different from T1 and T2, both p-values>.34). Thus, cortisol level increased after the stress manipulation and returned then back to the baseline level. There was no significant difference between the five measurements in the control group (all p-values>.52), thus showing no changes in cortisol level across the different testing sessions for participants not exposed to the stress manipulation.

To compute a reliable index of cortisol change due to the experimental manipulation, we considered the maximum value of the two samples collected right after the manipulation (T3 and T4) as *Post-manipulation* cortisol, and the minimum of the values referred to the samples outside the manipulation windows (before, T1 and T2, and at the end of the experiment, T5), as a measure of *Rest* cortisol. The differences in the salivary parameters in the *Post-Manipulation* and *Rest* cortisol, in the two groups, were compared with a two-way mixed ANOVA. Results show a main effect of *Group* (F(1,36)=12.50; p=0.001), a main effect of the *Manipulation* (Rest cortisol/ Post manipulation cortisol) (F(1,36)=33.50; p<0.001), and a significant interaction (F(1,36)=12.19; p=0.001). Post-hoc comparisons revealed that in the experimental group, the mean values of the salivary cortisol concentration of the post manipulation (M=0.531; SD=0.306) strongly increased as compared with the *Rest cortisol* values (M=0.191; SD=0.086; p<0.001). In the control group the *Post manipulation cortisol* (M=0.244; SD=0.155) did not differ from the baseline values (M=0.160; SD=0.056; p=0.25). The mean values at *Rest cortisol* were not different between the two groups p=0.62), whereas the experimental group showed higher cortisol levels than the control group in both post-manipulation measures (both p-values <0.001).

We then used these indices to quantify the magnitude of the changes in cortisol concentration induced by the experimental manipulation. For each participant, the values of the *Rest* cortisol were subtracted from those at the *Post-manipulation* cortisol to derive an index of *Cortisol Response* (CR). As shown in Fig.2A, while CR was small and homogenous in the control group (except for a single individual with CR>0.3 μg/dL), there was great variability in CR in the experimental group (see Fig.2B). Here, cortisol response in exhibited a bimodal distribution, with seven individuals showing high changes in CR, more than 0.3 μg/dL (Fig.2C), and the remaining participants showing CR changes smaller than 0.3 μg/dL. On this basis, we considered a threshold at <0.3 μg/dL as an index of cortisol response to the stress and we accordingly divided the experimental group into two sub-groups, the *“C-Responders”* and the *“C-Non-responders”* (CR<0.3 μg/dL) group.

**Figure 2.**
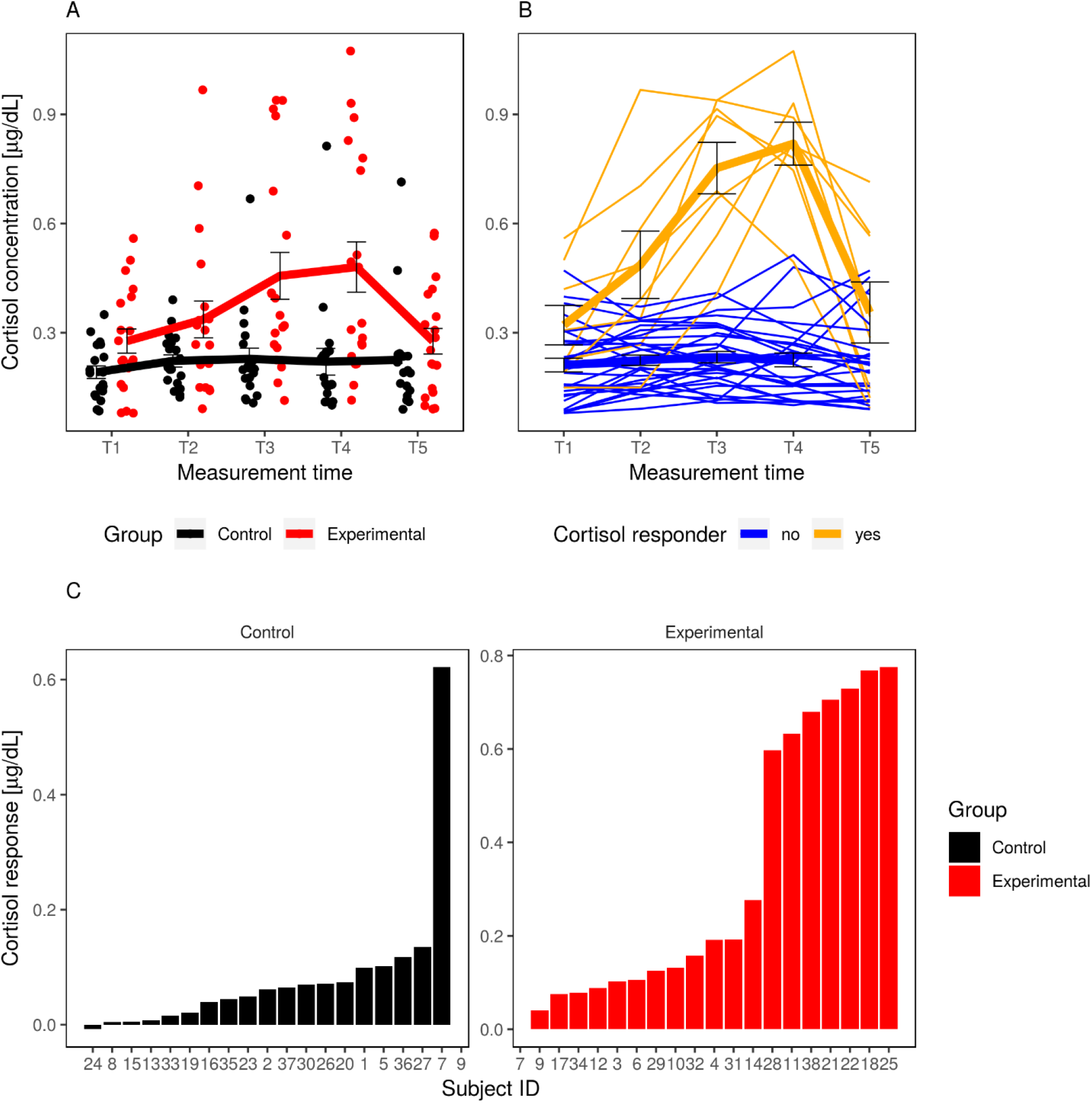
Cortisol response. (A). Individual salivary cortisol concentration expressed in μg/dL across the five Time-points for the Experimental (red) and Control (black) participants; bold lines indicate the group average, error bars represent S.E.M. (B) Individual trend of cortisol concentration of the experimental group, orange lines are representing the C-Responders, blue lines, the Non-C-Responders; bold lines indicate the group average, error bars represent S.E.M. (C) Bar plots representing the individual CR values of the cortisol concentration in the Control and the Experimental group. Values are obtained, for each participant, from the difference between the values of the Post-manipulation cortisol concentration (T3-T5) and the Rest cortisol concentration (T1, T2, T5).

### Subjective stress

We then test whether the experimental manipulation and the associated induced change in cortisol concentrations were reflected at the subjective level, measured with a 5-points Likert scale. Mean values of the subjective stress ratings across time were compared for the Experimental and the Control groups via non-parametric Friedman tests with the factor *Time-points (*T1, T2, T3, T4, T5), as the scores were not normally distributed. Neither for the experimental nor the control group, subjective stress reports significantly varied across time points (X^2^(4,19)=7.85; p=.1; X^2^(4,19)=8.30; p=.08), respectively).

To quantify the magnitude of the changes in the perceived subjective stress, induced by the experimental manipulation, we considered the mean value of the two measures collected right after the manipulation (T3 and T4) as Post-manipulation Subjective Stress, and the mean of the values referred to the measures outside the manipulation windows (before, T1 and T2, and at the end of the experiment, T5), as a measure of Rest Subjective Stress. Then, for each participant, the values of the *Rest* Subjective Stress were subtracted from those at the *Post-manipulation* Subjective Stress, to derive an index of the *Subjective Stress Response* (SSR).

We tested a possible relation between the subjective stress ratings and the cortisol concentration. There was no correlation between CR values and changes in SSR (p=0.41). Thus, changes in stress level as induced by the experimental manipulation and found at the cortisol level were not reflected in subjective ratings. This possibility was expected, as discussed in the introduction.

### Stress and PPS representation

To test whether the implicit neuroendocrine stress response was related to changes in the relation between near and far space, we analysed the two key parameters describing individuals’ PPS, i.e. the central point of the sigmoidal function, as a marker of the extent of the PPS, and the slope of the linear function, as a marker of the amount of near-far segregation in PPS representation. RTs for individual distances and the fitted linear function for different experimental conditions are shown in Figure 3.

**Figure 3.**
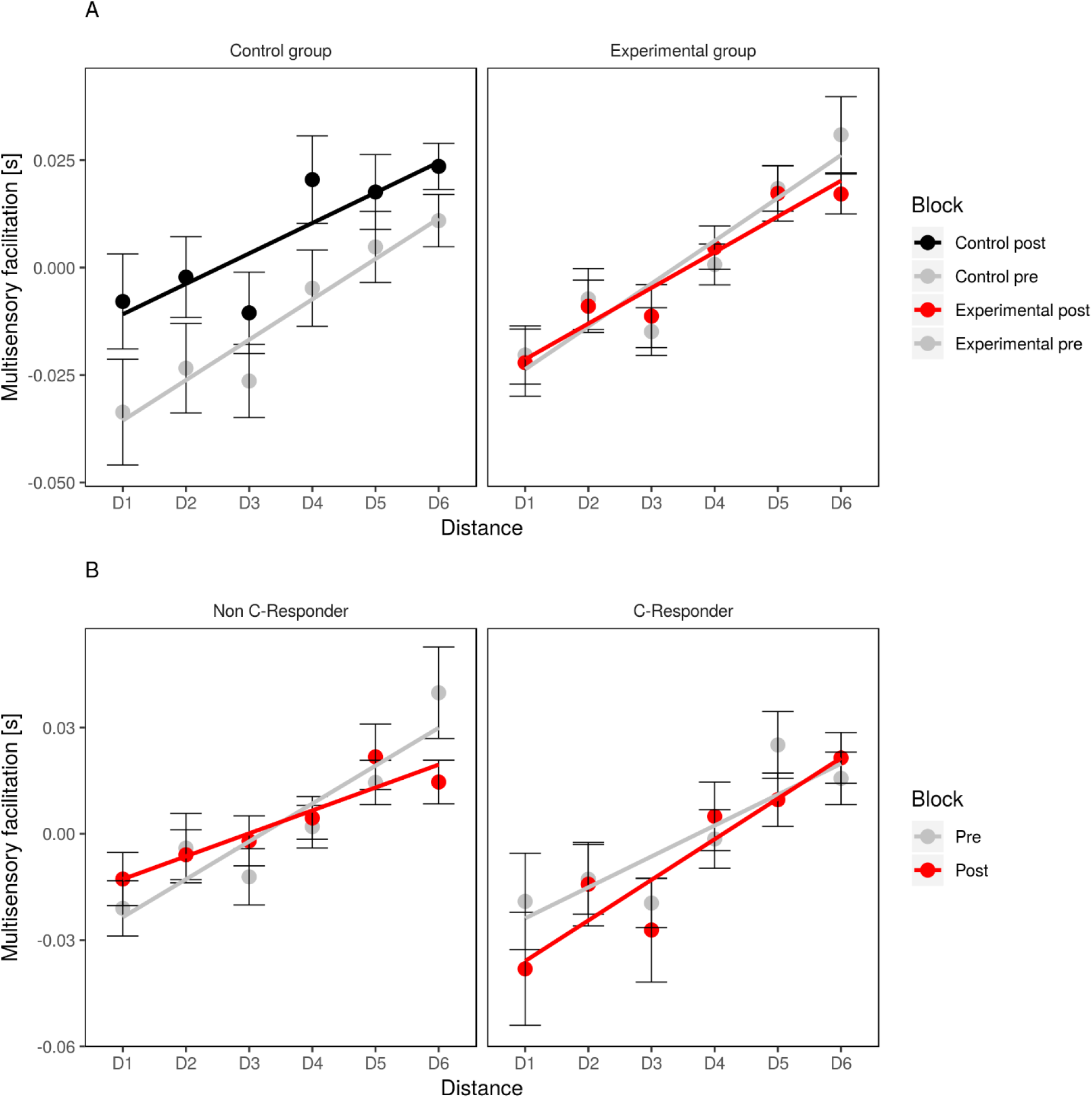
Peripersonal Space Results (RTs). (A). Multisensory facilitation (seconds) across distances, in the Pre and Post manipulation measurements. The left panel represents the results from the experimental group, the right panel, the results from the control group. The solid line represents the linear trend line (slope) of the distribution. Error bars represent S.E.M. (B). Multisensory facilitation (seconds) across distances, in the Pre and Post manipulation sessions, in the Non-C-Responders (left) and the C-Responders (right) sub-groups from the experimental group.

As a first general analysis, we compared changes in PPS representation, both for the central point and the slope, at the group level (the Experimental and the Control group), with two mixed ANOVAs with Group (Experimental/Control) and Manipulation (Pre/Post Manipulation). At the whole groups level, for the central point, there was no effect of Group, no effect of Manipulation, nor Interaction (all p-values>0.48). Similarly, no effects were found on the Slope (all p-values > 0.63). (Fig. 4A, right). The absence of interaction with the experimental group suggests that stressful manipulation does not change PPS representation at the group level.

**Figure 4.**
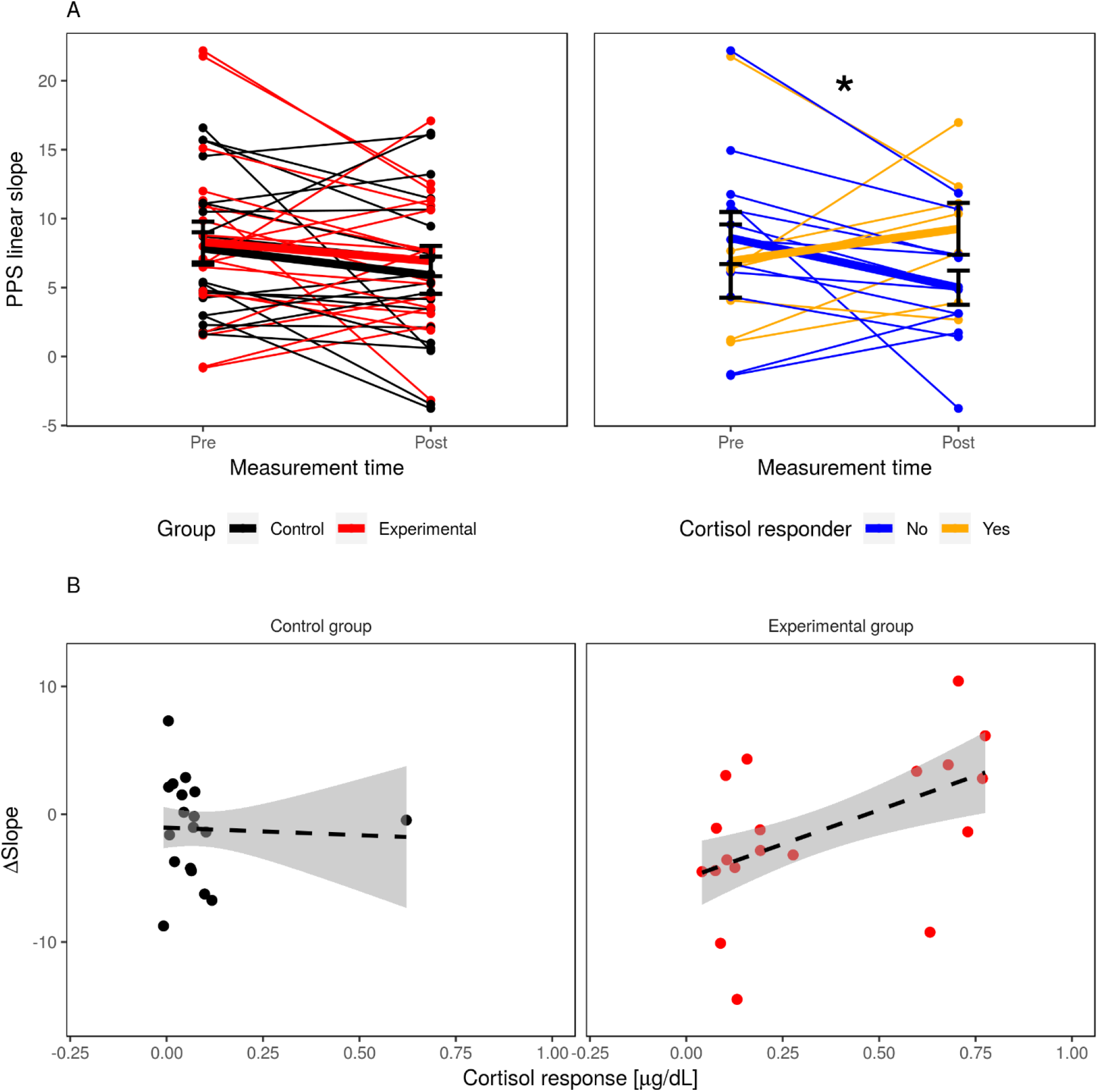
Peripersonal Space Results at the individual level (linear slopes). (A) In the left panel, PPS linear slopes for individual participants in the control (black) and experimental (red) group, shown before and after the manipulation. In the right panel, PPS linear slopes for the experimental group, split between C-Responders (orange) and Non-C-Responders (blue). Solid lines represent group means and their error bars represent S.E.M. (B). Scatterplot showing the relationship between Cortisol Response (μg/dL) and a difference in the PPS slope before and after the stressor manipulation (Δslope) in the Control group (left panel) and in the Experimental group (right panel). The dashed line represents the linear regression line. The more the participants were showing an increase in the cortisol response, the steeper the slope was after the stressor manipulation.

However, as seen from the analyses of cortisol concentration, the stress manipulation did not affect equally participants from the experimental group. Thus, to study the effect of induced physiological changes, we compared PPS representation between cortisol responders and nonresponders in the experimental group. To this aim, two separate ANOVAs were performed on the central point and on the slope of the fitted function with Cortisol Response (C-Responders and Non-C-Responders) as between-subjects factor, and Manipulation (Pre/Post Manipulation) as within-subjects factor. For the central point, the ANOVA showed no effects (all p-values>.13). Instead, for the Slope, we found a significant two-way interaction (F(1,17)=4.93; p=.040). Visual inspection of the pattern of multisensory slopes across the two experimental sessions shows that slopes in the C-Responders group increased after the manipulation, while it decreased in the Non-C-Responders group (Fig. 4A, left). In the pre-manipulation session no difference was found between the C-Responders and Non-C-Responders (C-Responders: M = 7.27, SD = 6.87, Non-C-Responders: Pre: M = 8.91, SD = 6.39, t(12) = 0.51, p = .62), while a trend toward significance was found in the post-manipulation session (C-Responders: M = 9.56, SD = 4.86, Non-C-Responders: M = 5.39, SD = 4.19, t(12) = −1.89, p = .084). Direct comparison between pre-post sessions in C-Responders and Non-Responders, showed a significant decrease in the slope for the Non-C-Responders (Pre: M = 0.891, SD = 0.639, Post: M = 0.539, SD = 0.419, t(11) = −2.41, p = .034), and an opposite direction, yet non-significant effect for C-Responders (Pre: M = 0.727, SD = 0.687, Post: M = 0.956, SD = 0.486, t(6) = 0.973, p = .36).

We then compared the effect of the Manipulation of the Non-C-Responders group with that of the Control group. We found a significant main effect of Manipulation (Pre: M = 8.25, SD = 5.57, Post: M = 5.70, SD = 5.22, F(1,29) = 8.27, p = .007), which did not interact with the Group (p = .41). The decrease of slopes in the Non-C-Responders group is comparable to the decrease observed in the control group (Pre: M = 7.83, SD = 5.13, Post: M = 5.89, SD = 5.88, t(18) = −1.627, p = .12), possibly reflecting a test-retest effect.

To further interpret the observed effect, we computed the difference in multisensory slope before and after the manipulation (*Δslope=Slope_Post_ – Slope_Pre_*) and fit a linear regression to predict the change of PPS slope as a function of the cortisol response, treated as a continuous variable:

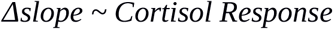

The model was significant (β = 0.507, R^2^ = 0.258, F(1,17) = 5.91, *p* = .026) and the positive coefficient indicates that participants with a stronger cortisol response had a larger increase in the multisensory slope, i.e.: in the amount of segregation between close and far space (Fig. 4B). The same regression fit on the control group yields no significant result (p = .71).

Furthermore, we tested whether an explicit measure of the stress response, detected by the *SSR* index, was related to changes in the PPS representation, measured on the slope and the central point values. Dividing both the control and the experimental group by the median values of the SSR, we obtained a *“low SSR”* group and a *“high SSR”* group. No changes in sigmoidal central point or linear fit slope between pre and post manipulation were highlighted in PPS representation by two-way ANOVAs, either for the control or the experimental group (all p-values > 0.12).

### Trait anxiety

As complementary analyses, we investigated the relationship between trait anxiety and neuroendocrine stress responses. Importantly, the two groups were not different from each other for the Trait-Anxiety Scores (Control: M=29.84, SD=8.62, N=19; Experimental: M=31.11, SD=5.56, N=19; p=0.59). Furthermore, we found no difference in trait anxiety between the sub-groups of *“C-Responder”* (M=30.75, SD=4.41, N=12) and *“Non-C-Responder”* (M=31.71, SD=7.499, N=7) (p=0.72).

However, we found a relationship between anxiety and stress, both at the subjective and physiological level. Trait-Anxiety Scores correlated with the Subjective Stress scores, both at the Baseline (*Control* group: R^2^=0.52, p=0.0005; *Experimental* group: R^2^=0.43, p=0.002) and the Post-Manipulation level for both groups (*Control* group: R^2^=0.60, p<0.001; *Experimental* group: R^2^=0.18, p=0.034). Trait Anxiety also correlated with cortisol level at *Baseline* (R^2^=0.169; p=0.011). However, anxiety scores did not predict changes in cortisol (*CR* values) induced by the experimental manipulations (*Control* group: R^2^=0.0001, p=0.96; Experimental group: R^2^=0.01, p=0.66).

Finally, we investigated whether our main results were also influenced by state anxiety. We found no relation between state anxiety scores and PPS slope in the pre-manipulation block (p=0.24, control and experimental group together), nor with the change in PPS slope in the experimental group (p=0.73). When using a multiple regression predicting the change in slope with state anxiety scores and cortisol responses, only the latter was found to have a significant effect, as in our main analyses (cortisol response: p=0.03; state anxiety: p=0.52).

## Discussion

PPS representation can be mainly conceived as a multimodal sensory-motor interface which mediates the interaction between the body and external objects by integrating information about external stimuli (i.e., visual stimuli) with body-related cues (i.e., tactile stimuli) to prime appropriate reactions (Graziano and Cooke, 2006; Làdavas and Serino, 2008; Cléry et al, 2015). In the present study, we tested whether acute stress, eliciting a significant neuroendocrine response, affects PPS representation, measured by changes in the processing of body-related multisensory stimuli in space. Importantly, given the significant individual differences in stress responsiveness described in the literature, we tested whether the magnitude of the cortisol level increase determined the changes in PPS representation. Thus, we distinguished two subgroups in the participants exposed to the stressor manipulation, accordingly to whether there was a sensible and meaningful change in their neuroendocrine response, *C-Responders* or *Non-C-Responders*.

At the group level, we did not find a general PPS change in the Experimental group as compared to the control group consequent to the stressor manipulation. However, the changes in PPS representation, associated with the manipulation, were different among individuals who did, or did not, show a neuroendocrine stress response after stress exposure. In the experimental group, the PPS linear slope increased in participants who showed a cortisol response to the stressful manipulation, whereas it decreased in participants who did not have such a response. The decrease in slope in Non-C-Responders was comparable to what observed in the control group, suggesting this effect can be due to habituation to the experimental task. The central point of the sigmoidal fit was not affected by the manipulation either, indicating that the stress response does not affect the extent of PPS. Moreover, subjects from the control group, who were not exposed to a stressful situation, did not show any difference in PPS representation measured before and after the manipulation, neither in terms of slope nor of central point. Taken together, this evidence suggests that stress affects PPS representation in terms of a change in the way external stimuli in space interact with the processing of tactile information on the body. Namely, individuals presenting a significant physiological stress response - i.e., *C-Responders* - showed an enhanced differentiation between the close and the far space after the stressful manipulation. This effect is in line with a recent study by Spacassassi and Maravita (2020), demonstrating, with a multisensory temporal order task, an enhanced multisensory interaction for stimuli presented in the near as compared to the far space, after an anxiety-inducing manipulation. Importantly, the change was significant only when considering individual differences in the sensitivity to the manipulation and anxiety levels.

The steeper slopes of the PPS in *C-Responders* reflect an increase of multisensory processing within a more limited area around their body, and thus a stronger differentiation between near and far space compared with the baseline. We propose that such an increase in PPS segregation might be considered as a form of defensive-freezing reaction, whereby resources are more allocated to the body and space immediately surrounding it (Schmidt et al, 2008; Roelofs, 2017). It is worth reminding that “freezing” should not be confused with a state of hypoactivity (Hagenaars, Oitzl and Roelofs, 2014), but it has been rather described as a state of “attentive immobility” (Roelofs, 2017). This defensive response serves evolutionary advantages in optimizing the selection of an appropriate coping response by enhancing perceptual and attentional processes (Lang, Davis and Öhman, 2000; Erickson, Drevets and Schulkin, 2003), that become more automatic and less controlled (Arnsten, 2009; Sänger et al, 2014; Elling et al, 2012) and in action preparation (Gladwin et al, 2016). We propose that the enhanced segregation of PPS, i.e., preferential integration for near space in stress *C-Responders* might reflect “freezing” at the PPS level. In this respect, it is worth noting that previous studies showed an increased PPS linear slope in the presence of looming stimuli, specifically when considered relevant, as in the case of a threat (de Haan et al, 2016).

There is a complex interaction between motor responses, PPS representation and stress reaction. To interact with external objects, the individual has to be able to predict the spatio-temporal relationship between an external stimulus and one’s own body (Bourgeois and Coello, 2012; Pfeiffer et al, 2018; Noel et al., 2015; Van Elk, Forget, and Blanke, 2013; Cardinali, Brozzoli, and Farnè, 2009; Serino et al, 2011; Graziano, Reiss, and Gross, 1999). Multisensory integration within the PPS is a key mechanism underlying such prediction (Clèry et al, 2015), and its immediate translation in a potential action in a dynamic environment, via modulation of the motor system (Finisguerra et al, 2015; Makin et al, 2009; Serino et al, 2009). Results have shown that PPS dynamically shapes according to the experience of controlling the course of events through one’s actions (D’Angelo et al, 2018), according to the characteristics of external stimuli (e.g., velocity; Noel et al, 2018), previous exposure to the specific movement (Brozzoli, Gentile, and Ehrsson, 2012; Brozzoli et al, 2011; Noel et al, 2015) and the dimension of the acting space (Bassolino et al, 2014; Canzoneri, Ubaldi, et al, 2013). Conversely, previous studies in mice models have highlighted that an active control over a stressor, or the possibility to act, modulate the dynamics of the stress response (Fox, Merali, and Harrison, 2006; Kunz, 2014). Rodents that underwent uncontrollable stressful situations for a prolonged time showed less inhibited behaviour and less depressive traits when they could act over the stressor, as compared with the individuals constrained in a passive condition (Laborit, 1976, 1988). Interestingly, within the domain of the social-spatial representations, Iachini and colleagues (2014) demonstrated that, when participants have active control over social interaction, they are less sensitive to a confederate intruding on their comfort space. Similarly, the increased allocation of multisensor-motor resources to the far space in cortisol non-responders, as opposed to responders, may indicate a tendency to actively cope with the stressful situation.

In our study, the relationship between responsiveness to stress and changes in PPS representation was limited to the physiological measures of stress, as we did not find any difference in participants showing a higher increase of subjective stress as compared to those showing a lower increase. However, subjective stress ratings were generally low, their change between baseline and post-manipulation assessments did not even distinguish between the control and the experimental group, and finally, they were not related to cortisol response. Thus, the Likert scale used to assess subjective stress was possibly not sensitive enough to capture changes induced by our manipulation. Alternatively, the physiological cortisol response does truly relate to a different component of the stress phenomenon.

Finally, to identify predictors of responsiveness to stress, we also collected and analysed trait anxiety scores from the present sample. First, although anxiety scores were correlated with the level of stress at the baseline, both for subjective and cortisol measures, they were not related to the stress changes induced by the manipulation for either measure. Also, anxiety scores were not correlated with the indices of PPS representation, unlike in previous studies by Sambo and Iannetti (2013) and Iachini and colleagues (2015), who found that higher levels of anxiety were associated with a more extended PPS, as to increase monitoring of potential threats. This difference might depend on the nature of the PPS task implemented. Here, stimuli were more neutral, while, in the above-cited studies, stimuli were highly arousing (eye-blink reflex induced by median nerve electric shock, or intrusion by an estranger into one’ own comfort space). Thus, a relationship between anxiety and PPS might emerge if more salient or arousing stimuli are involved.

To conclude, here we report a novel relation between responsiveness to stress, as measured at the physiological level by cortisol concentrations, and a change in multisensory integration of stimuli in space, following acute exposure to stress. We showed an increased differentiation between the multisensory processing of near vs. far stimuli in participants showing a physiologically measurable stress response, which might reflect a freezing-like response at the PPS level. Given the role of PPS representation in processing multisensory stimuli to prepare appropriate motor responses, the present results suggest that stress reflects in a re-allocation of multisensory-motor resources in the space immediately surrounding the body. Therefore, these findings could provide useful insights on how an easily obtainable biometric measure, such as cortisol concentration, could predict individuals’ capability to monitor the multisensory space and act in a stressful situation, allowing them to better define and counteract its negative impact.

## Limitations

A possible limitation of the present study should be acknowledged. In our design, the effect of stress was to be investigated by comparing the experimental group with a control group. However, the effect of the experimental manipulation was best explained by considering interindividual differences in the stress response within the experimental group, reducing the effective sample size to the experimental group only. Further investigations should address this limitation by directly using a within-subjects design, allowing to maximise statistical power.

## Acknowledgements

This research was funded by SNSF Professorship Grant to Andrea Serino (PP00P3_163951). The authors thank Olivia Zanoletti for performing the cortisol analyses, Elisa Pellencin for pilot data collection, Javier Belo-Ruiz, Robin Mange, Bruno Herbelin and Olaf Blanke for using the RealiSM technology.

## Author contributions

Giulia Ellena: Formal analysis, Writing-Original Draft, Visualization. Tommaso Bertoni: Formal analysis, Writing-Original Draft, Visualization. Manon Durand-Ruel: Conceptualization, Methodology, Investigation. John Thoresen: Conceptualization, Methodology, Investigation. Carmen Sandi: Conceptualization, Methodology, Resources, Writing - Review and Editing, Supervision. Andrea Serino: Conceptualization, Methodology, Resources, Writing - Review and Editing, Supervision.

